# Trait-environment interactions influence tree planting success in Peruvian tropical dry forests

**DOI:** 10.1101/2025.02.21.639418

**Authors:** W. Goossens, T. Fremout, B. Muys, J.C. Soto Quispe, K. Van Meerbeek, S.L. Maes

## Abstract

Tropical dry forests (TDFs) are among the most endangered ecosystems globally, despite their critical socio-ecological importance. In recent decades, tree-planting initiatives have been widely implemented to restore these forests, yielding mixed results. To enhance restoration effectiveness, it is imperative to adopt a predictive approach, focusing on the drivers of seedling performance and their interactions. This study explores how functional traits of planted tree species interact with site environmental conditions to influence seedling survival and growth rates across nine restoration sites of varying ages and climatic conditions within the Peruvian Tumbes-Piura TDFs. While the widely accepted resource-use theory predicts that acquisitive species – focusing on the fast retrieval of resources and characterised by higher specific leaf area and lower wood density – achieve high growth rates in resource-rich environments but experience sharp declines under resource-limited conditions, we found that acquisitive species maintained their high growth rates along the environmental gradient. In contrast, conservative species, expected to be less sensitive to resource variability, exhibited marked declines in growth when water or nutrient availability decreased. Additionally, nitrogen-fixing species did not outperform non-fixers in growth or survival, indicating that nitrogen fixation alone does not confer a consistent advantage in early restoration. Overall, our findings highlight the importance of trait-environment interactions in tree planting performance and underscore the need to align species selection and management strategies with site conditions and broader restoration goals to enhance the success of TDF restoration.

**Implications for practice:**

- Our findings highlight the importance of considering trait-environment interactions when evaluating growth rates across TDF restoration sites.
- Planting acquisitive species, which maintained higher growth rates than conservative species even under resource-poor environmental conditions, can accelerate tropical dry forest recovery in resource-poor restoration sites within the Peruvian Tumbes-Piura region.
- While seedling growth is strongly shaped by environmental factors and their interaction with functional traits, survival may depend more on site-specific factors and management interventions, including irrigation and fertilisation.
- Selecting species based solely on nitrogen fixation ability may not enhance restoration success, as nitrogen-fixing species did not exhibit higher growth or survival than non-fixers.

## INTRODUCTION

All around the globe, forests face significant threats from deforestation and degradation, leading to biodiversity loss, accelerating climate change and negatively impacting human well-being (FAO & UNEP 2020). In response to these alarming trends, there has been a surge in ambitious commitments to restore degraded forest ecosystems across all continents (Sewell et al. 2020), further amplified by the current ‘UN Decade on Ecosystem Restoration’ (2021-2030), which underscores the urgency of reviving damaged ecosystems (United Nations General Assembly 2019).

Many restoration initiatives involve the active planting of trees to speed up ecosystem recovery (Brancalion & Holl 2020; Mesa-Sierra et al. 2024), which is particularly important in tropical dry forests (TDFs) that experience a dry season of several months (Mooney et al. 1995). Active planting can considerably accelerate forest establishment in these areas, as natural recovery is often slow and uncertain due to harsh microclimates and severe forest fragmentation (Alvarez-Aquino & Williams-Linera 2012; Dimson & Gillespie 2020).

Due to numerous limitations, TDFs remain among the most threatened terrestrial ecosystems (Miles et al. 2006), despite their socio-ecological importance in housing numerous endemic species and supporting the livelihoods of some of the world’s poorest communities (Blackie et al. 2014; Schröder et al. 2021). The urgency to restore these ecosystems has led to various TDF restoration initiatives (Campo et al. 2023; Dimson & Gillespie 2020; Cárdenas et al. 2022). However, tree planting as TDF restoration approach often yields variable and disappointing growth and survival rates (Dimson & Gillespie 2020; Werden et al. 2023), despite substantial resource investments (Toro et al. 2024). Restoration ecology is therefore urged to become more predictive by studying the factors influencing seedling performance regarding growth and survival (Campo et al. 2023; Brudvig & Catano 2021; Lanuza et al. 2020). Both the species’ *functional identity* and the site’s *environmental conditions* play a crucial role in governing growth and survival rates of planted trees (Rosell et al. 2022; Werden et al. 2020). Hence, understanding how both factors relate to each other is key to tailoring restoration designs successfully to local environmental contexts and improving overall restoration success (Eviner & Hawkes 2008; Zirbel & Brudvig 2020; Werden et al. 2023).

The functioning of tree species can be summarised by placing them along the *acquisitive-*

*conservative* functional trait spectrum (Díaz et al. 2004; Wright et al. 2010). By grouping species with similar resource-use strategies, we can identify and predict their response to environmental gradients (Pinho et al. 2019) and adapt management practices accordingly (Eviner & Hawkes 2008). In TDFs, interactions between species’ functional traits and environmental conditions have been explored to understand patterns of functional diversity along edaphic, climatic and topographic gradients (Méndez-Toribio et al. 2017; Comita & Engelbrecht 2009; Pinho et al. 2019), suggesting that trait-environment interactions play an important role in shaping TDF tree growth and survival rates (Sterck et al. 2011).

TDF species have developed distinct resource-use strategies to cope with the seasonally limited water availability, which is widely regarded as the main limiting factor for seedling growth and survival in these forests (Alvarez-Aquino & Williams-Linera 2012; Méndez-Toribio et al. 2020; Markesteijn & Poorter 2009). On the one hand, drought-avoidant acquisitive species are usually deciduous and characterised by a rapid growth response to changes in water availability (Rosell et al. 2022). They possess relatively large transpirational tissues (i.e. high specific leaf area (SLA)) and low wood density (WD), facilitating greater hydraulic conductivity during the wet season (Poorter et al. 2019; Sterck et al. 2011; Matlaga et al. 2024). In contrast, drought-tolerant conservative species, with higher wood density and lower SLA, prioritise resource retention and structural reinforcement, resulting in slower growth but enhanced stress tolerance and survival (Werden et al. 2018).

General resource-use theory predicts that acquisitive species perform better under resource-rich, favourable environmental conditions, but experience sharp performance declines when resources become scarce (Méndez-Toribio et al. 2020; Worthy et al. 2020; Mira et al. 2023). Indeed, their high growth rate drops rapidly when water (Sterck et al. 2011) or soil nutrient (Baribault et al. 2012; Reich 2014) availability decline. Low soil water availability or high evaporative demand increases the water potential difference between soil and atmosphere (Bauman et al. 2022; Méndez-Toribio et al. 2020), forcing acquisitive species to close their stomata to prevent hydraulic failure, resulting in lower growth rates and potentially even carbon-starvation-induced mortality after prolonged drought (McDowell et al. 2008). Conservative species are less sensitive to hydraulic failures and can maintain photosynthetic activity under drier conditions (Sterck et al. 2011; Matlaga et al. 2024). Regarding soil nutrient limitations, acquisitive species require higher soil fertility, giving them a growth advantage in fertile soils but posing challenges when nutrients are scarce (Worthy et al. 2020; Reich 2014). Moreover, while some studies suggest that acquisitive species are more sensitive to rising temperatures than conservative species (Sastry & Barua 2017), others contradict this (Slot & Winter 2017).

An additional important trait in TDFs is nitrogen fixation, which can provide a competitive advantage through increased photosynthetic activity and water-use efficiency (Pellegrini et al. 2016; Menge & Chazdon 2016). As nitrogen-fixing species span the full acquisitive-conservative spectrum (Gei 2014), both resource-use strategy and nitrogen fixation should be considered in tropical forest analyses (Batterman et al. 2018; Werden et al. 2023).

Since seedling performance, i.e. their growth and survival rates, is influenced by both species traits (acquisitive-conservative spectrum) and local environmental contexts (resources and conditions), their interactions might play a pivotal role in restoration practices (Caleño-Ruiz et al. 2023; Zirbel & Brudvig 2020; Mira et al. 2023) and explain some of the observed variability in restoration outcomes (Dimson & Gillespie 2020). However, the trait-performance relationship is not always consistent, with some studies showing no clear link between resource-use strategies and tree performance (Martínez-Garza et al. 2013; Poorter et al. 2018). Additionally, while research on trait-environment has expanded, most studies have focussed on tropical rainforests (Matlaga et al. 2024; Worthy et al. 2020), neglecting the TDF biome (Campo et al. 2023; Schröder et al. 2021). This study aims to improve understanding of how functional trait identity and environmental conditions interactively influence seedling performance across nine TDF restoration projects. We focus on the Tumbes-Piura dry forest region in northwestern Peru, contributing to improved tree-planting practices in the area (Cerrón et al. 2019; Fremout et al. 2022).

Specifically, we evaluate the impacts of interactions between abiotic (i.e. climate and soil) conditions and functional trait identities (i.e. principal component axis of WD and SLA) of ten adult tree species on the performance (i.e. growth and survival) of planted seedlings across nine real-world restoration sites covering a gradient of environmental conditions. We expect that acquisitive species (with lower WD and higher SLA) will (1) experience lower mortality and higher growth rates in sites with higher fertility and lower atmospheric aridity, but (2) be more sensitive than conservative species to lower resource availabilities (Sterck et al. 2011; Zirbel & Brudvig 2020). Thus, we anticipate conservative species to show little variation in seedling performance across the studied environmental gradient, whereas acquisitive species are expected to show higher mortality and lower growth rates in unfavourable conditions but to exhibit increased performance as environmental conditions improve (Matlaga et al. 2024).

## MATERIALS AND METHODS

### Study sites and region

To evaluate the drivers of tree planting performance, we sampled nine restoration sites in the lowland Tumbes-Piura dry forests in northwestern Peru (Figure 1), situated between the Pacific coast and the Andean mountain range up to altitudes of 500 m.a.s.l. (Linares-Palomino & Alvarez 2005). The forests host 234 species, with 6 to 25 species per hectare, reaching a basal area of 2.31 to 22.79 m²/ha (Linares-Palomino & Alvarez 2005; Linares-Palomino et al. 2010). Mature forests in the region typically reach a canopy height of 10 to 15 metres. Average annual precipitation in the Tumbes-Piura region is low, ranging between 100 and 500 mm, of which around 85% is centred between January and April, coinciding with the warmest period of the year (Best & Kessler 1995; Cueva-Ortiz et al. 2020). Mean annual temperatures vary between 20 and 26°C (Cueva-Ortiz et al. 2020). Every 3 to 7 years, the El Niño Southern Oscillation (ENSO) brings more intense rains to the region, with precipitation levels up to ten times higher than the climatic mean, fostering higher (natural) tree recruitment and growth rates (Best & Kessler 1995).

**Figure 1:**
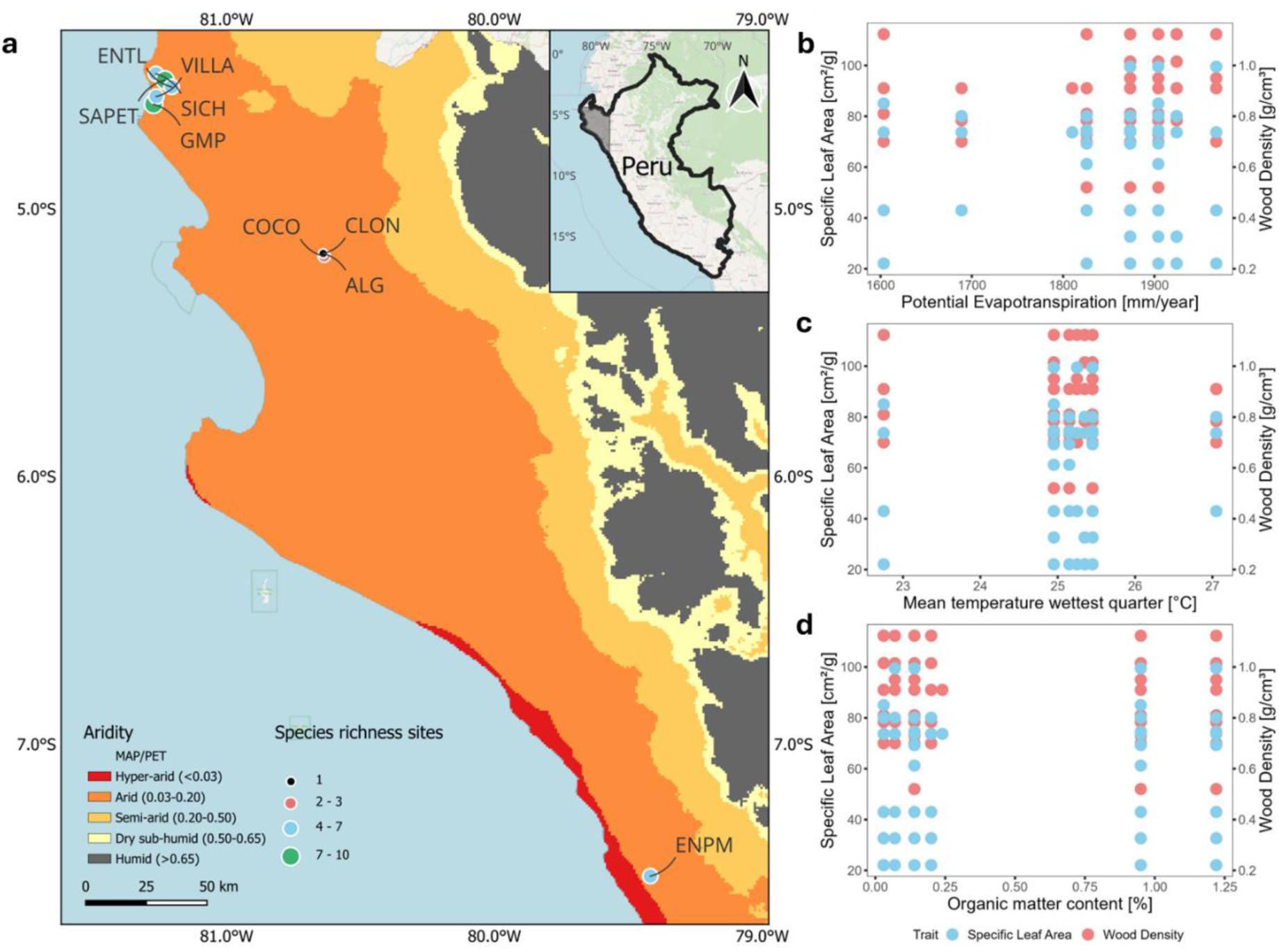
(a) Map of the aridity index (mean annual precipitation divided by potential evapotranspiration (MAP/PET) following Zomer et al. (2022)) showing study site locations and the number of planted species at each site across northwestern Peru. Distribution of functional traits – specific leaf area [cm²/g] and wood density [g/cm³] (see further) – along gradients of (b) potential evapotranspiration [mm/year], (c) mean temperature of the wettest quarter [°C], and (d) organic matter content [%]. Map generated using QGIS 3.34.0 (QGIS Development Team 2024), graphs created with R 4.2.3 (R Core Team 2023).

The Tumbes-Piura dry forests belong to the Tumbes-Chocó-Magdalena biodiversity hotspot (CEPF 2005) and are renowned for their endemism (Best & Kessler 1995; Linares-Palomino et al. 2010). Nonetheless, substantial portions of these forests have been lost through deforestation, and much of the remaining patches are strongly degraded due to overgrazing by livestock and logging (Fremout et al. 2020; Linares-Palomino & Alvarez 2005). In recent years, the region has witnessed many restoration efforts to reverse this trend, aiming to restore forest cover and structure, conserve biodiversity, and raise awareness among the local population (Cerrón et al. 2019). Between August and September 2022, we sampled nine tree-planting restoration sites in the area, varying in size (0.22 - 12.7 ha), planting year (1988 – 2022), and species composition (Supplement S1). Three suitable sites were obtained from the project database of Cerrón et al. (2019) and complemented with six projects suggested by restoration managers in the region. The sites covered a gradient in climatic conditions (Figure 1), including potential evapotranspiration (1603 - 1968 mm/year), mean annual precipitation (63 – 188 mm) and mean temperature of the wettest quarter (22.75 – 27.05 °C). Although we obtained some data on restoration design – planting density (200 – 4444 individuals/hectare) and propagation method (bare-rooted – containerised) across eight sites – and management practices – irrigation (40 – 80 L/plant/month) and fertilisation (1.2 – 2.5 kg (organic) fertiliser/plant/year) at four sites – this information was incomplete and hence omitted from further analyses (Supplement S1).

### Abiotic conditions

Both climatic and soil conditions were assessed for their influence on planted seedling performance (Supplement S1). Since direct quantification of some resources available to seedlings – such as water and energy availability – is challenging (Parkin et al. 2006), we selected established environmental conditions affecting resource availability. We extracted climate variables for every site from the CHELSA 2.1 Climate Database (Karger et al. 2017) for the period 1981 to 2010 in QGIS 3.34.0 (QGIS Development Team 2024). These variables, with a resolution of 30 arcsec (ca. 0.9 km at the equator), included mean annual temperature (MAT), mean daily air temperatures of the wettest (BIO8) and driest (BIO9) quarter, mean annual precipitation (MAP) and mean annual potential evapotranspiration (PET), computed by summing up monthly averages. Rather than directly quantifying resource availability, these variables describe environmental conditions regulating water and energy balance, with MAT, BIO8 and BIO9 reflecting thermal conditions (Denissen et al. 2022), and MAP and PET capturing aspects of water availability (Konapala et al. 2020; Li et al. 2022; Stefanidis & Alexandridis 2021). For example, PET, indicating atmospheric aridity, has previously explained differing performances of conservative and acquisitive species in TDFs (Méndez-Toribio et al. 2020).

To account for soil variability, eight subsamples were collected per site and combined using the quartering method (EPA 2023) to produce one representative composite sample (Pennock et al. 2006). The subsamples were taken at depths of 10, 20 and 30 cm from eight evenly spaced locations in a zigzag pattern. For two sites exhibiting clear soil heterogeneity, multiple composite samples were acquired from different zones to assess within-site variance, resulting in four composite samples (i.e. 4 × 8 subsamples) at site ENTL and two at site SAPET. The composite soil samples were then air-dried and stored in plastic bags until analysis at the *Universidad Nacional Agraria La Molina* for organic matter content, electrical conductivity, pH and concentrations of K, P and CaCO_3_ (Supplement S1).

### Functional traits

To evaluate the effect of functional traits on seedling performance, we included WD, SLA, maximum height (H), seed mass (SM), deciduousness and nitrogen fixation (Supplement S2).

WD [g/cm³] and SLA [cm²/g] were measured from wood cores and leaf samples for five species (i.e. *Neltuma pallida, Beautempsia avicenniifolia, Parkinsonia aculeata, Lycium boerhaviifolium* and *Morisonia scabrida*), while for *Loxopterygium huasango* only SLA was obtained in the field, as we did not receive permission to extract wood cores for the site in which the species was present as an adult. The SLA and WD values of the other four species (i.e. *Cordia lutea*, *Libidibia glabrata*, *Tara spinosa* and *Vachellia macracantha*) were obtained from literature, as we could not find healthy, mature trees to sample from. Similarly, the maximum tree height, seed mass, deciduousness and nitrogen fixation ability of all studied species were obtained from literature.

SLA was determined by collecting ten undamaged leaves fully exposed to the sun from two to four isolated mature individuals near the restoration sites (Cornelissen et al. 2003; Pérez-Harguindeguy et al. 2013). We included the leaf petioles and did not subdivide compound leaves into individual leaflets. Fresh leaf area was calculated in the software program ImageJ (Schneider et al. 2012) following the protocol of O’Neal et al. (2002). After oven-drying the leaves for 48 hours at 80 °C and measuring their mass, the SLA was calculated by dividing the leaf area by the oven-dry weight. For the wood density measurements, we sampled mature individuals in the vicinity of the planting sites with a tree increment borer (Chave 2006). The extracted cores were stored in paper straws, dried at 80°C for 48 hours (Cornelissen et al. 2003), and the density was computed using X-ray Computed Tomography in the Woodlab at Ghent University in Belgium (Van den Bulcke et al. 2014). The scanned images were converted to mean densities of the whole core using a reference material of known density and air with a density assumed to be zero.

After standardising the continuous traits (SM, H, WD and SLA), we conducted a principal component analysis (PCA) with the *prcomp* function (R Core Team 2023) and retrieved the first two axes explaining 65% of the trait variation (Jollife & Cadima 2016). Similar to Díaz et al. (2015), we found that one axis corresponded to the resource-use strategy of species (PC_resource-use_), with higher values corresponding to species with a lower SLA and higher WD (Supplement S2). The other axis (PC_size_) was strongly associated with H and SM, increasing when species showed higher maximum height and lower seed mass.

### Tree growth and survival

We measured the height and recorded survival status of every seedling across the nine study sites (Moonlight et al. 2021). For multi-stemmed trees, commonly found in TDFs, only the tallest stem was recorded. Survival status was evaluated by scratching the bark and observing whether the cambium displayed signs of life (Werden et al. 2020). Relative height growth rates (growth rates [ln(m)/year]) were computed by subtracting the natural logarithm of the obtained height by the natural logarithm of 0.30 m, reported as the mean height at planting by five restoration managers, and divided by the time in years between planting and monitoring (Werden et al. 2023; Worthy et al. 2020):

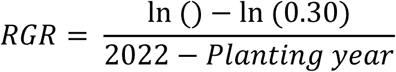

During the survival assessment, we could only evaluate the status of individuals present at the time of monitoring for the seven mixed-species sites, which meant that empty planting spots had to be omitted, leading to an overestimation of survival rates and potentially introducing bias by favouring some species more than others. However, in the two oldest sites, which are monocultures of *Neltuma pallida*, species identity was not a concern, allowing accurate quantification of mortality rates.

### Data analysis

Overall, ten species were present in at least two sites and were included in our analysis. For the two sites established in the monitoring year, we only evaluated survivorship, as measured heights likely represent planting height, and not (yet) significant growth rates due to the short growing time period (Rodríguez-Alarcón et al. 2024). We removed all 21 RGR observations considered outliers at the site level following Tukey (1977). Our final dataset consisted of 5514 tree individuals for the survival assessment, which resulted in 3057 growth measurements after removing dead individuals, growth outliers and the two sites established in the monitoring year (Supplement S3). All explanatory factors, except for the categorical nitrogen fixation and two PCA-axes, were standardised to facilitate model fitting and effect size comparison (Zuur et al. 2009).

Generalised linear mixed models (GLMMs) were employed to evaluate the drivers of growth and survival variability in TDF restoration sites, with site identity as a random intercept to account for the ecological assumption that within-site observations are more closely related (Banin et al. 2023), while species identity was also included as a random factor. In sites with multiple composite soil samples, tree individuals were related to their soil sampling zone. The model included the first two PCA-axes, depicting the variation in resource-use (PC_resource-use_) and size (PC_size_) among the species (Díaz et al. 2015; Liu et al. 2022; Werden et al. 2023), and ability to fix nitrogen (Nfix). We expanded this three-factor model with the environmental factors potential evapotranspiration (PET), mean temperature in the wettest quarter (BIO8) and presence of organic matter (OM). Given the strong correlations between the environmental factors, only a limited set was used as explanatory variables. We included PET as a proxy of water availability, considering that it indicates atmospheric aridity and has been linked previously to differing growth responses of conservative and acquisitive species (Méndez-Toribio et al. 2020). Similarly, interactive effects are expected between resource-use strategy and BIO8 – an indicator for energy availability in the rainy season (Slot & Winter 2017; Denissen et al. 2022) – and OM – representing the soil fertility and water-holding capacity (Wood et al. 2016). Interaction effects with PC_resource-use_ were included for all abiotic predictors except BIO8, due to induced multicollinearity and increased variance inflation factors (VIFs) (Kim 2019). Moreover, we included site age as a fixed variable in the growth model, accounting for time-related changes in microclimate and growth patterns (Werden et al. 2020; Eviner & Hawkes 2008). While age is also relevant to model survivorship (Liu et al. 2022), its strong correlation with the site-level random effect caused model singularity. Since age’s effect was indirectly captured by the site random effect in the survival model, the following two models were ultimately tested:

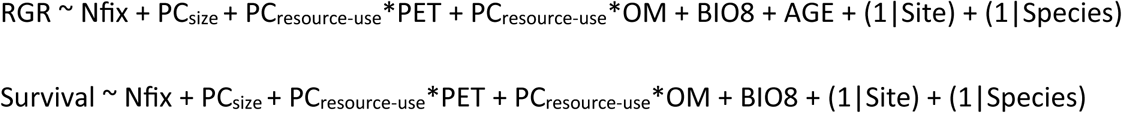

Height growth rates were analysed using the *lme4* package (Bates et al. 2015) with Gaussian error distribution and Restricted Maximum Likelihood (REML) estimation (Banin et al. 2023; Zuur et al. 2009). Significance of the results was assessed via Satterwaite’s degrees of freedom method in the *lmerTest* package (Kuznetsova et al. 2017). Survival was modelled with a binomial error distribution and *cloglog* link function (Banin et al. 2023; Liu et al. 2022) using the *glmmTMB* package (Brooks et al. 2017). We used the *piecewiseSEM* package (Lefcheck 2016) to compute the marginal (R²_m_) and conditional (R²_c_) coefficients of determination to assess the goodness-of-fit (Nakagawa & Schielzeth 2013). The former quantifies the variance explained by the fixed effects, while the latter evaluates the variance explained by both the fixed and random factors. Model assumptions were checked using the *DHARMa* package (Hartig 2024). Two additional models were tested, following the same structure but including the interaction between resource-use strategy and BIO8 instead of PET. We also ran the survival model including age as an additional explanatory variable and omitting the random effect of site (Supplement S4), which confirmed the importance of age in explaining survival rates. All analyses were performed in R 4.2.3 (R Core Team 2023), using the *tidyverse* (Wickham et al. 2019), *ggeffects* (Lüdecke 2025) and *ggpubr* (Kassambara 2023) packages.

## RESULTS

Tree seedling growth varied considerably across species and sites (Supplement S3) and was significantly influenced by trait-environment interactions (Table 1; Figure 2a-b). First, the interaction between PC_resource-use_ and PET interactively influenced height growth. Specifically, the growth of conservative species – with a higher WD and lower SLA – was more negatively affected by an increase in PET than that of acquisitive species (PET × PC_resource-use_ −0.09 p < 0.001). Likewise, a significant interaction existed between PC_resource-use_ and OM, as the height growth of acquisitive species remained relatively stable and even slightly increased with lower values of OM, while that of conservative species decreased when OM decreased (OM × PC_resource-use_ 0.10 p < 0.001). Significant main effects were found for BIO8 (0.48 p < 0.05) and age (0.04 p <0.05), while nitrogen fixation and PC_size_ had no significant impact. Overall, the fixed effects accounted for 75.3% of the variance in height growth (R²_m_), increasing to 82.3% when site and species were included as random factors (R²_c_).

**Table 1:** Parameter estimates (β) and 95%-confidence intervals (CI) for the generalised linear mixed models (GLMMs) examining height growth and survival rates focusing on the interaction between resource-use strategy and water availability. Predictor variables include nitrogen fixation (Nfix), resource-use strategy (PC_resource-use_), size variation (PC_size_), potential evapotranspiration (PET [mm/year]), organic matter content (OM [%]), mean temperature in the wettest quarter (BIO8 [°C]) and site age. Low values of PC_resource-use_ indicate acquisitive species, while higher values correspond to more conservative species. Significance of the parameters is indicated with * p<0.05, ** p<0.01 and *** p<0.001. Goodness-of-fit is given as both marginal (R²_m_) and conditional (R²c) coefficients of determination. Supplement S6 provides additional details on the degrees of freedom and t-values for the growth model, and z-values for the survival model.

| Driver | Height growth rate |  | Survival rate |  |
| --- | --- | --- | --- | --- |
| | $\beta$ | CI | $\beta$ | CI |
| Nfix | -0.17 | [-0.36, 0.02] | 0.48 | [0.06, 0.90] |
| $PC_{\text{resource-use}}$ | -0.29** | [-0.35, -0.23] | -0.11 | [-0.25, 0.04] |
| $PC_{\text{size}}$ | 0.16 | [0.07, 0.24] | -0.11 | [-0.30, 0.09] |
| PET | -0.31* | [-0.40, -0.23] | 0.05 | [-0.11, 0.21] |
| OM | 0.10* | [0.06, 0.14] | -0.04 | [-0.08, -0.01] |
| BIO8 | 0.48* | [0.38, 0.57] | -0.25 | [-0.41, -0.09] |
| AGE | 0.04* | [0.04, 0.05] | NA | NA |
| $PC_{\text{resource-use}} \times PET$ | -0.09*** | [-0.11, -0.08] | -0.03 | [-0.06, 0.00] |
| $PC_{\text{resource-use}} \times OM$ | 0.10*** | [0.07, 0.12] | 0.06* | [0.03, 0.09] |
| $R^2_m$ (%) | 75.3 | | 4.6 | |
| $R^2_c$ (%) | 82.3 | | 24.6 | |

**Figure 2:**
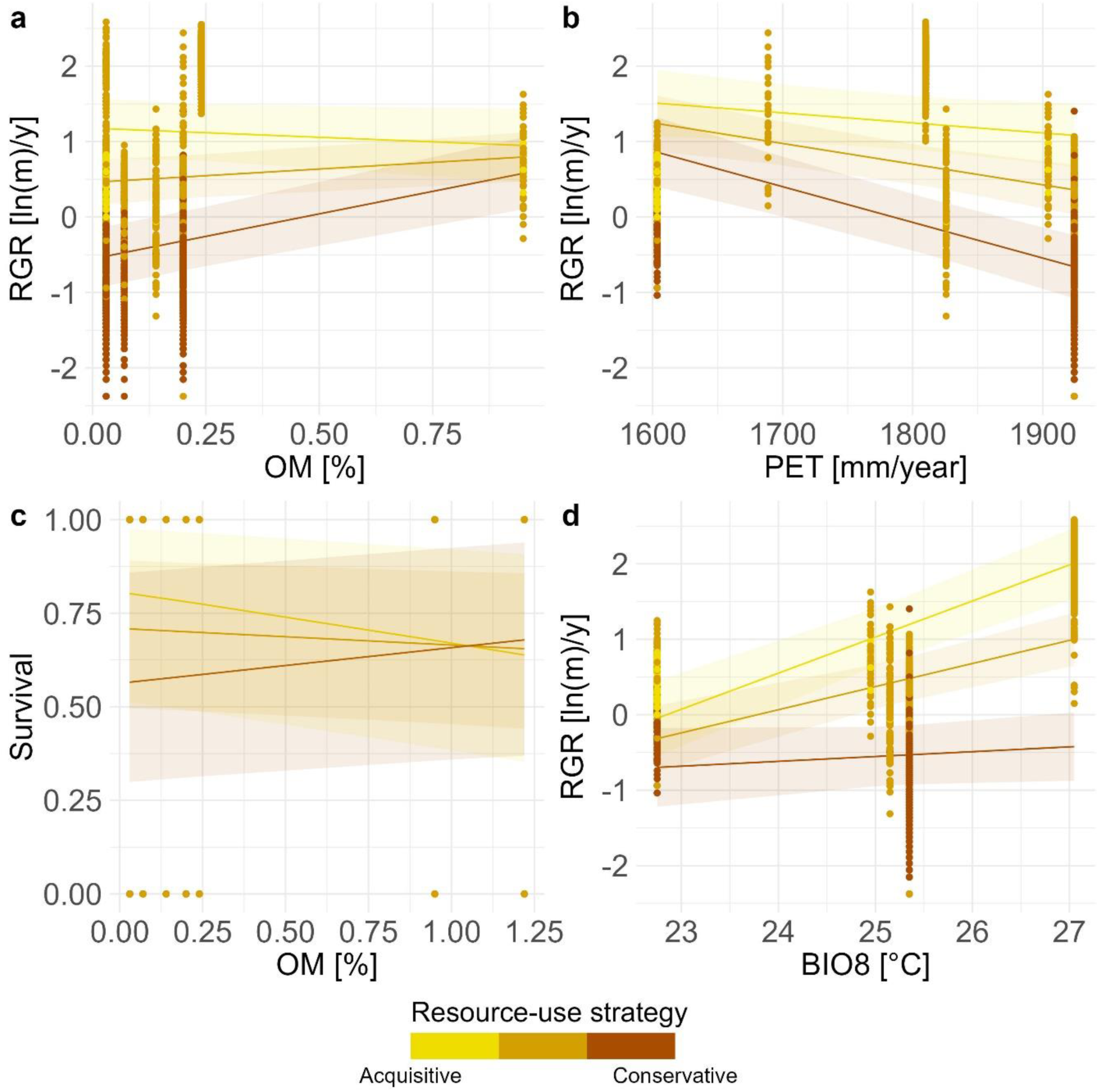
Significant interaction effects from the first model between resource-use strategy (acquisitive versus conservative based on the first principal component axis) and (a) annual potential evapotranspiration (PET [mm/year]) and (b) organic matter content (OM [%]) on height growth. Effect (c) of the interaction between resource-use strategy and organic matter content on survival rates. Results from the additional model, (d) showing the significant interaction effect between the resource-use strategy and mean temperature during the wettest quarter (BIO8 [°C]) on height growth. Lines represent predicted values for three levels of the first principal component axis, corresponding to the most acquisitive, an intermediate and the most conservative species. Points represent individuals, coloured according to their resource-use strategy.

In contrast, while seedling survivorship also varied considerably across species and sites (Supplement S3), the only significant effect in the survival model was a positive interaction between OM and PC_resource-use_ (OM × PC_resource-use_ 0.06 p < 0.05). Specifically, survival rates of conservative species increased with higher organic matter content, but the opposite trend was observed for acquisitive species (Table 1; Figure 2c). However, the fixed effects could only explain 4.6% of the variation (R²_m_) in the survival model, increasing to 24.6% when accounting for the random intercepts of species and site (R²_c_).

When testing an alternative model replacing the PET interaction with an interaction between BIO8 and PC_resource-use_, results remained largely consistent. BIO8 and PC_resource-use_ significantly interacted to influence height growth (Figure 2d), with acquisitive species showing stronger growth responses to higher temperature in the wettest quarter than conservative species (BIO8 × PC_resource-use_ 0.16 p < 0.001). The OM × PC_resource-use_ interaction remained significant (OM × PC_resource-use_ 0.09 p < 0.001), showing the same pattern as in the PET interaction model. Similarly, the survival model including the interaction with BIO8 instead of PET yielded comparable results, with the OM × PC_resource-use_ interaction remaining the only significant effect (OM × PC_resource-use_ 0.07 p < 0.05). Given these consistencies, we present only the BIO8 interaction for growth in Figure 2, with additional statistical details for the growth and survival models in Supplement S5.

## DISCUSSION

Our results demonstrate the importance of incorporating interactions between functional traits and environmental variables when assessing tree planting performance in restoration projects. While most studies assess the individual effects of these factors on seedling growth and survival (Werden et al. 2018; Rosell et al. 2022), the presence of significant interactions suggests that considering them in isolation may not fully capture the complexity of TDF restoration projects.

### Acquisitive species are less sensitive to higher evapotranspiration and lower organic matter content

Acquisitive species – characterised by lower WD and higher SLA – maintained relative high growth rates across the environmental gradients in potential evapotranspiration (as a proxy for atmospheric aridity and water availability; Méndez-Toribio et al. 2020) and organic matter content (a proxy for nutrient content; Wood et al. 2016). This suggests that acquisitive species were able to retrieve sufficient resources and allocate them to height growth across a range of resource availability conditions, i.e. even under resource-poorer conditions. This finding opposes the general resource-use theory, which predicts higher growth rates for acquisitive species under favourable conditions but a sharp decline in performance when water and nutrients become limiting, while, conversely, conservative species are expected to be less sensitive to such resource variation (Sterck et al. 2011; Baribault et al. 2012; Caleño-Ruiz et al. 2023).

The growth rates of acquisitive species appeared less sensitive to the environmental variation (i.e. both water and organic matter availability), which indicates that once established, acquisitive seedlings can sustain high growth rates regardless of site resource availability, suggesting a competitive growth advantage over slow-growing conservative species (Werden et al. 2023). Mechanistically, a possible explanation is that fast-growing acquisitive species invest heavily in belowground biomass and thereby increase their rooting density and depth (Fagundes et al. 2022; Méndez-Toribio et al. 2020; Martínez-Garza et al. 2013), which enhances their growth capacity (Werden et al. 2023) and reduces susceptibility to changes in soil fertility and atmospheric aridity (Markesteijn & Poorter 2009). Once well-established, acquisitive species may thus retrieve nutrients and water more efficiently than conservative species, allowing them to sustain height growth (Carrasco-Carballido et al. 2019). However, due to the lack of root trait data, we cannot confirm this hypothesis, although previous studies support a link between acquisitive resource use and rooting density (Comita & Engelbrecht 2009; Méndez-Toribio et al. 2020).

Another possible explanation for the relatively sustained growth rates of acquisitive species along the resource gradient is that they prioritise high growth rates during the short wet season in TDFs (Mendivelso et al. 2014; Poorter et al. 2018), after which many quickly shed their leaves to avoid drought stress (Fagundes et al. 2022; Poorter & Markesteijn 2008; Rodríguez-Alarcón et al. 2024). Conservative species, in contrast, remain active during (part of) the dry season, extending their growing period but also increasing their susceptibility to the higher evaporative demand (McDowell et al. 2008; Wright et al. 2010). As a result, conservative species are forced to adjust their growth rates to match available resources, making them more sensitive to increased potential evapotranspiration.

Our survival analysis further revealed contrasting effects of organic matter availability, with higher nutrient content increasing survivorship in conservative species but reducing it in acquisitive species. Although previous studies reported higher survival rates for fast-growing acquisitive species in early TDF restoration stages (Werden et al. 2023; Martínez-Garza et al. 2013; González-Tokman et al. 2018), the observed decline at greater nutrient availability is unexpected. However, given the low explanatory power of our survival model, these results should be interpreted with caution.

### Acquisitive species benefit more from higher wet season temperatures

Higher temperatures during the wettest quarter positively affected growth rates, challenging our initial hypothesis and previous studies (Méndez-Toribio et al. 2017; Mendivelso et al. 2014). One possible explanation is that increased temperatures during the growing season enhance energy availability, thereby stimulating photosynthetic activity (Slot & Winter 2017). We also anticipated that acquisitive species would be more sensitive than conservative species to temperature increases, expecting a more rapid growth decline under warmer conditions (Sastry & Barua 2017). Although we could not include this interaction in the main model due to high VIFs, testing it separately revealed that acquisitive species exhibited a stronger positive response to temperature than conservative species. This supports the hypothesis by Slot and Winter (2017) that acquisitive species benefit more from increased energy availability, likely due to their higher electron transport capacity and greater plasticity in temperature responses. Additionally, these results align with our hypothesis that acquisitive species are less sensitive to higher atmospheric aridity, as they concentrate growth during the wet period when water and energy availability are optimal.

### Nitrogen fixation and size do not significantly affect seedling performance

Contrary to our hypothesis, nitrogen-fixing species did not exhibit enhanced growth and survival rates, despite potentially higher photosynthetic activity and water-use efficiency (Pellegrini et al. 2016; Menge & Chazdon 2016). As in previous TDF studies (Werden et al. 2023; Cárdenas et al. 2022), nitrogen-fixing and non-fixing species showed similar performance, suggesting that nitrogen fixation alone does not necessarily provide an advantage in early restoration sites. This reinforces the importance of including mixtures of fixers and non-fixers in restoration projects (Werden et al. 2023), especially considering that nitrogen fixing species span the acquisitive-conservative resource-use gradient (Gei 2014).

Similarly, we found no significant effect of the principal component axis representing variation in species size. This axis was positively associated with maximum height but negatively with seed mass, whereas Díaz et al. (2015) found that both traits contributed positively to size variation. This suggests that trait relationships among our study species may differ from those observed in other systems, possibly explaining why the size axis was not a strong predictor of seedling performance, in contrast to previous studies on tropical forests that found important, although contrasting, effects of seed mass and maximum height on seedling growth and survival (Wright et al. 2010; Martínez-Garza et al. 2013).

### Study limitations and future research avenues

Overall, our results indicate that across nine restoration projects in the Tumbes-Piura TDFs, acquisitive species maintain relative high growth rates even when water or nutrient content becomes limiting, and respond positively to higher temperatures during the wet season. Additionally, the relatively small difference between the marginal and conditional R² values in the growth model suggests that the fixed main and interactive effects effectively captured growth variation. These findings highlight the importance of trait-environment interactions in seedling performance, providing valuable empirical evidence to inform tree planting strategies in TDF restoration.

However, we acknowledge the limitations caused by our study’s observational nature, which restricts our ability to directly assess key management strategies, initial planting conditions and seedling provenance (Dimson & Gillespie 2020; González-Tokman et al. 2018), and prevented control over factors like the planted number of individuals per species. Additionally, while temperature, potential evapotranspiration and organic matter content are useful proxies for resource availability (Stefanidis & Alexandridis 2021; Konapala et al. 2020; Li et al. 2022), they do not fully capture the complexity of water and energy dynamics (Parkin et al. 2006; Bauman et al. 2022) or soil fertility (Pennock et al. 2006). The scarcity of initial planting data also meant that variations in initial seedling sizes might have influenced the relative growth rate calculations, although much of this variation is likely captured by the random effects of species and site. Similarly, we could only include remaining dead individuals with certain species identification and had to omit empty planting spots in all but two sites, which may have resulted in an overrepresentation of conservative species with more durable stems in the survival models (Chave et al. 2009), potentially biasing survival estimates and contributing to the models’ low explanatory power. Notably, the relatively high variance in survivorship explained by random effects indicates that unmeasured species- and site-specific factors, including local management interventions (Mesa-Sierra et al. 2024), may have influenced survival rates.

Nonetheless, our study provides important insights into resource-use dynamics within real-world tree planting efforts in TDFs, complementing findings from controlled but relatively small-scale experiments (Werden et al. 2020, 2023) and broader observational studies of natural forest succession (Poorter et al. 2019; Derroire et al. 2018). While growth was largely shaped by trait-environment interactions, our findings suggest that survival may depend more strongly on factors such as management practices (Dimson & Gillespie 2020; Mesa-Sierra et al. 2024). In contrast, Toro et al. (2024) found limited effects of irrigation and fertilisation on the growth and survival of most TDF species. However, their findings may not be directly comparable to ours, as their study was conducted in a region with substantially higher mean annual precipitation and shorter dry season.

Future research on trait-environment interactions should prioritise controlled experimental designs with known initial species numbers, management practices and environmental conditions along a gradient in both water and nutrient availability. We also encourage future restoration research to incorporate trait plasticity (Derroire et al. 2018; Poorter et al. 2018) and ontogenetic variability (Metz et al. 2023; Poorter & Markesteijn 2008), since both factors allow individuals to acclimate to local abiotic conditions and could govern trait-environment interactions in restoration projects (Lanuza et al. 2020; Worthy et al. 2020). Additionally, accounting for the high soil and microclimatic heterogeneity of TDFs in future studies can shed additional light on the importance of trait-environment interactions in restoration practices (Rodríguez-Alarcón et al. 2024; Caleño-Ruiz et al. 2023).

Given our observation that seedling growth depends on the interaction between environmental variables and species’ resource-use strategy, our findings underscore the importance of context-dependent management approaches in restoration projects and highlight that species selection should consider both the site’s environmental conditions and overarching restoration objectives (Zirbel & Brudvig 2020; Fremout et al. 2022), while also integrating management interventions to enhance seedling survival (Brancalion & Holl 2020). This study lays the foundation for future research on these interactions, particularly within the underexplored Tumbes-Piura TDFs.

## Acknowledgements

The authors express their gratitude to Michael Perring (UKCEH) for his valuable critique of the manuscript and to Nora Grados and Gaston Cruz (UDEP) for their assistance with legal documentation and the analysis of leaf and wood samples. The authors also acknowledge the cooperation of restoration managers who facilitated the research on their sites, as well as the Peruvian Forest Service (SERFOR) for granting research and botanical sampling permissions. The fieldwork was made possible through a VLIR-UOS scholarship (REI-2022-01-01).

