## Supplement S1 for "Trait-environment interactions influence tree planting success in Peruvian tropical dry forests"

### Supplement S1: Site characteristics

Tables i and ii give a complete overview of available site characteristics and extracted environmental factors characterising our sites. Soil samples were obtained in the field and analysed at the *Universidad Nacional Agraria La Molina* ([*http://www.lamolina.edu.pe/facultad/agricola/lasmaf_servicios.htm*](http://www.lamolina.edu.pe/facultad/agricola/lasmaf_servicios.htm)), while for the climatic factors the CHELSA 2.1 Climate Database was used (Karger et al. 2017). Data on the management practices were obtained from the restoration managers. Site names are abbreviations used in the field and have been anonymised for privacy reasons.

Table i: Overview of site size [ha], number and identity of the species, and number of individuals per hectare (Density [Individuals/ha]). Known management practices included are: seedling propagation method (bare-rooted or containerised seedlings), protection against herbivores (yes or no), irrigation [L/plant/month] and fertilisation [kg/plant/year]. In case only qualitative data was available, the variable is included in a binary way (yes or no), while unavailable data is denoted as NA.

| **Site** | **Size [ha]** | | **Species  number** | **Age  [year]** | **Density [Ind/ha]** | **Planting approach** | **Herbivore  protection** | **Irrigation [L/plant/month]** | **Fertilisation [kg/plant/year]** | **Species [number of individuals]** |
| --- | --- | --- | --- | --- | --- | --- | --- | --- | --- | --- |
| ALG | | 4.58 | 1 | 34 | 400 | Bare-rooted | Yes | Yes | NA | *N. pallida* [861] |
| CLON | | 1.25 | 1 | 23 | 400 | Bare-rooted | Yes | Yes | NA | *N. pallida* [151] |
| COCO | | 0.22 | 3 | 6 | 400 | Containerised | Yes | Yes | Yes | *L. boerhaviifolium* [7]*, N. pallida* [18]*, P. aculeata* [4] |
| ENPM | | 2.13 | 4 | 6 | 400 | Bare-rooted | No | 42 | 2.5 | *M. scabrida* [71]*, N. pallida* [67]*, P. aculeata* [33]*,*  *T. spinosa* [111] |
| ENTL | | 12.7 | 4 | 8 | 200 | Bare-rooted | No | 40 | 2 | *B. avicenniifolia* [119]*, L. boerhaviifolium* [51],  *M. scabrida* [1064]*, N. pallida* [1381] |
| GMP | | 0.39 | 10 | 12 | NA | NA | NA | NA | NA | *B. avicenniifolia* [30]*, C. lutea* [10]*,*  *L. boerhaviifolium* [22], *L. glabrata* [7]*,*  *L. huasango* [1]*, M. scabrida* [2]*, N. pallida* [30]*,*  *P. aculeata* [7]*, T. spinosa* [11]*, V. macracantha* [1] |
| SAPET | | 5.7 | 8 | 0 | 278 | Containerised | Yes | 80 | 1.2 | *B. avicenniifolia* [46]*, C. lutea* [14]*, L. glabrata* [2]*,*  *L. boerhaviifolium* [14]*, M. scabrida* [33]*,*  *N. pallida* [538]*, P. aculeata* [3]*, V. macracantha* [49] |
| SICH | | 2.51 | 4 | 0 | 278 | Containerised | Yes | 80 | 1.2 | *L. glabrata* [2]*, M. scabrida* [1]*, N. pallida* [579]*,*  *P. aculeata* [1] |
| VILLA | | 0.17 | 7 | 3 | 4444 | Containerised | Yes | Yes | Yes | *C. lutea* [3]*, L. huasango* [25]*,*  *L. boerhaviifolium* [136]*, M. scabrida* [1]*,*  *N. pallida* [12]*, P. aculeata* [1]*, V. macracantha* [4] |

*Table ii: Abiotic factors describing the site conditions, with MAT being the mean annual temperature [°C], BIO8 [°C] and BIO9 [°C] the temperature in the wettest respectively driest quarter, MAP the mean annual precipitation [mm/year], PET the annual potential evapotranspiration [mm/year] and Aridity the aridity index (MAP/PET). Elev gives the mean elevation of the site [m]. Soil variables include pH, electrical conductivity (EC [dS/m]) and the presence of calcium-carbonate (CaCO3 [%]), organic matter (OM [%]), phosphorus (P [ppm]) and potassium (K [ppm]). For the ENTL and SAPET sites, four respectively two soil samples were obtained.*

| **Site** | **MAP [°C]** | **BIO8 [°C]** | **BIO9 [°C]** | **MAT [mm/year]** | **PET [mm/year]** | **Arid** | **Elev [m]** | **Age [year]** | **pH** | **CE [dS/m]** | **CaCO3 [%]** | **MO [%]** | **P [ppm]** | **K [ppm]** |
| --- | --- | --- | --- | --- | --- | --- | --- | --- | --- | --- | --- | --- | --- | --- |
| ALG | 24.05 | 27.05 | 21.45 | 188 | 1810 | 0.1039 | 41 | 34 | 7.68 | 0.75 | 0.09 | 0.24 | 5.70 | 208.00 |
| CLON | 24.05 | 27.05 | 21.45 | 188 | 1810 | 0.1039 | 41 | 23 | 8.17 | 0.59 | 0.27 | 0.03 | 3.90 | 158.00 |
| COCO | 24.05 | 27.05 | 21.45 | 187 | 1689 | 0.1105 | 37 | 6 | 7.86 | 5.72 | 1.79 | 0.20 | 2.50 | 258.00 |
| ENPM | 19.75 | 22.75 | 17.75 | 63 | 1603 | 0.0389 | 110 | 6 | 8.36 | 4.56 | 2.24 | 0.03 | 1.80 | 126.00 |
| ENTL | 22.55 | 25.35 | 20.15 | 188 | 1924 | 0.0953 | 98 | 8 | 7.59 | 11.69 | 0.00 | 0.03 | 7.30 | 253.00 |
|  |  |  |  |  |  |  |  |  | 7.69 | 16.82 | 0.00 | 0.03 | 2.80 | 235.00 |
|  |  |  |  |  |  |  |  |  | 7.55 | 12.73 | 0.00 | 0.07 | 4.50 | 189.00 |
|  |  |  |  |  |  |  |  |  | 7.64 | 6.43 | 0.00 | 0.20 | 4.10 | 191.00 |
| GMP | 22.05 | 24.95 | 19.75 | 149 | 1904 | 0.0796 | 78 | 12 | 7.77 | 12.13 | 5.28 | 0.95 | 4.90 | 314.00 |
| SAPET | 22.55 | 25.45 | 20.15 | 172 | 1873 | 0.0915 | 22 | 0 | 7.96 | 1.64 | 0.00 | 1.22 | 4.80 | 202.00 |
|  |  |  |  |  |  |  |  |  | 8.22 | 0.82 | 0.45 | 0.14 | 3.80 | 129.00 |
| SICH | 22.35 | 25.25 | 20.05 | 162 | 1968 | 0.0824 | 13 | 0 | 8.40 | 0.76 | 1.34 | 0.07 | 3.10 | 126.00 |
| VILLA | 22.15 | 25.15 | 19.85 | 157 | 1826 | 0.086 | 56 | 3 | 7.55 | 25.46 | 12.07 | 0.14 | 3.90 | 224.00 |

*Measurement techniques:

pH: measured with a potentiometer in a soil: water suspension of 1:1

EC: electrical conductivity of the aqueous soil: water extract at rate of 1:1

CaCO3: gas-volumetric method using a calcimeter

OM: Walkley and Black method, oxidation of organic carbon with potassium dichromate. %OM = %C × 1.724

P: Modified Olsen method, extraction with NaHCO₃ 0.5M, pH 8.5.

K: Extraction with ammonium acetate (CH₃ - COONH₄)N, pH 7.0.

### Supplement S2: Functional traits and principal component analysis

Here we give an overview of functional trait measurements and literature values corresponding to the respective species (Table iii-v). Note that for multiple species limited or no data are available in the literature, and genus mean values can differ greatly from actual values. Consequently, we chose to keep our measured functional traits in the analysis. Figure i shows the first two axes of the principal component analysis (PCA), with names adjusted following Díaz et al. (2015).

Table iii: List of analysed species and their measured wood densities (in bold) with standard deviation and the number of samples across the number of sites between parentheses. Literature values are added, together with their reference and if applicable the method of calculation (note).

| **Scientific name** | **WD  [g/cm³]** | **WD literature [g/cm³]** | **Note** | **References** |
| --- | --- | --- | --- | --- |
| *Beautempsia avicenniifolia* | **1.02 ± 0.12** (7 \| 2 sites) | 0.75 | Caparis mean | Álvarez et al. 2013 |
| *Cordia lutea* | 0.52 | 0.52 | Genus mean | Montalván 2015 |
| *Libidibia glabrata* | 0.95 | 0.95 |  | Zanne et al. 2009 |
| *Loxopterygium huasango* | 0.73 | 0.73 |  | Zanne et al. 2009 |
| *Lycium boerhaviifolium* | **0.78 ± 0.05** (3 \| 2 sites) | 0.64 | Genus mean | Choat et al. 2012 |
| *Morisonia scabrida* | **1.12 ± 0.09** (7 \| 3 sites) | 0.77 |  | Zanne et al. 2009 |
| *Neltuma pallida* | **0.91 ± 0.11** (14 \| 5 sites) | 0.88 |  | Zanne et al. 2009 |
| *Parkinsonia aculeata* | **0.70 ± 0.08** (5 \| 4 sites) | 0.67 | Tropical South America | Zanne et al. 2009 |
| *Tara spinosa* | 1.05 | 1.05 | Genus mean | FAO, 1997 |
| *Vachellia macracantha* | 0.81 | 0.81 |  | Montalván 2015 |

Table iv: List of analysed species and their measured specific leaf areas (in bold) with standard deviation and number of sites in which ten leaves were collected between parentheses. Literature values are added, together with their reference and if applicable the method of calculation. Note that we were not able to find SLA values for two species, i.e. Beautempsia avicenniifolia and Lycium boerhaviifolium.

| **Scientific name** | **SLA  [cm²/g]** | **SLA literature [cm²/g]** | **Note** | **References** |
| --- | --- | --- | --- | --- |
| *Beautempsia avicenniifolia* | **32.6 ± 1.42** (2) | NA |  |  |
| *Cordia lutea* | 74.60 | 74.60 |  | Montalván 2015 |
| *Libidibia glabrata* | 99.50 | 99.50 |  | Montalván 2015 |
| *Loxopterygium huasango* | **61.2** (1) | 137.70 |  | Montalván 2015 |
| *Lycium boerhaviifolium* | **80.1 ± 18.6** (2) | NA |  |  |
| *Morisonia scabrida* | **22.1 ± 1.37** (2) | 45.60 |  | Montalván 2015 |
| *Neltuma pallida* | **73.7 ± 11.9** (3) | 125.10 | Species mean 1/LMA | Salazar et al. 2018 |
| *Parkinsonia aculeata* | **32.3 ± 14.4** (2) | 39.10 | Species mean | Kabir et al. 2020 |
| *Tara spinosa* | 86.00 | 85.00 | Species mean |  |
| *Vachellia macracantha* | 69.30 | 69.30 |  | Montalván 2015 |

Table v: List of analysed species and their family name with the ability to fixate atmospheric nitrogen (N-Fixation: True (T) or False (F)), leaf habit (evergreen, semi-deciduous or deciduous), seed mass (SM [mg]) and maximum height (H [m]). Data on nitrogen fixation, deciduousness and maximal height were obtained from Fremout et al. (2023), references for the seed mass data are given in the last column.

| **Family** | **Scientific name** | **N- Fixation** | **Leaf**  **Habit** | **SM**  **[mg]** | **H [m]** | **References seed mass** |
| --- | --- | --- | --- | --- | --- | --- |
| Capparaceae | *Beautempsia avicenniifolia* | F | Evergreen | 91.22 | 2 | Montalván 2015 |
| Cordiaceae | *Cordia lutea* | F | Deciduous | 105.5 | 4 | Montalván 2015 |
| Fabaceae | *Libidibia glabrata* | T | (Semi-)  deciduous | 106.3 | 13 | Montalván 2015 |
| Anacardiaceae | *Loxopterygium huasango* | F | Deciduous | 7 | 25 | SID 2025 |
| Solanaceae | *Lycium boerhaviifolium* | F | Evergreen | 1.95 | 3 | Mora-Costillo et al. 2020 |
| Capparaceae | *Morisonia scabrida* | F | Evergreen | 33.3 | 10 | Montalván 2015 |
| Fabaceae | *Neltuma pallida* | T | (Semi-)  deciduous | 38.7 | 15 | SID 2025 |
| Fabaceae | *Parkinsonia aculeata* | F | Evergreen | 105 | 12 | SID 2025 |
| Fabaceae | *Tara spinosa* | F | Evergreen | 228.49 | 15 | SID 2025 |
| Fabaceae | *Vachellia macracantha* | T | (Semi-)  deciduous | 31.2 | 12 | Montalván 2015 |

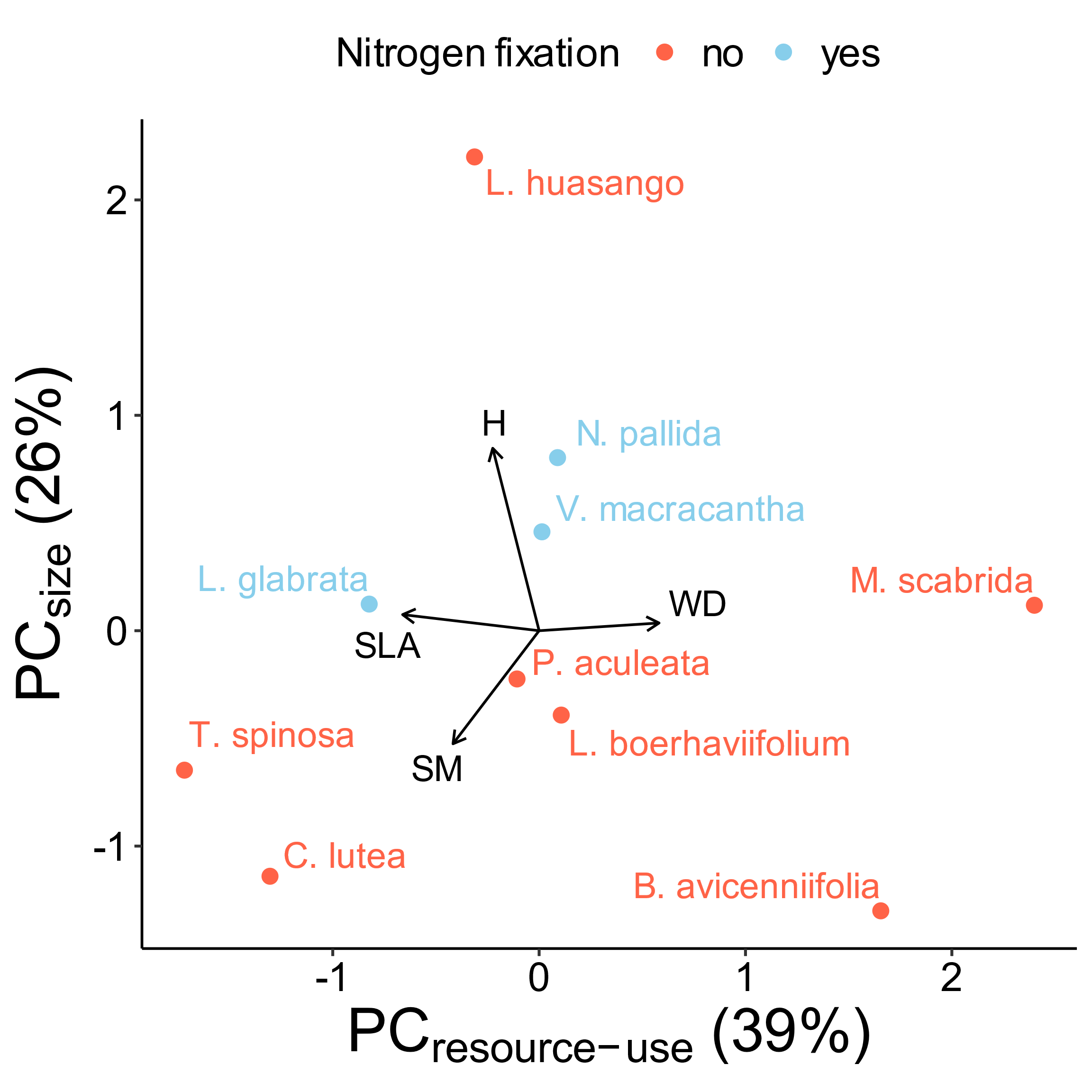

Figure i: Biplot of principal component analysis of tree species trait variation. The first axis explained 39% of the variation and was mainly influenced by WD and SLA, while the second axis accounted for 26% of trait variation among species, characterised by the effects of H and SM. Axes names were adjusted according to the literature. Nitrogen-fixing species are denoted in blue, while non-fixing species are coloured red.

### Supplement S3: Performance data

An overview of the growth (RGR) and survival rates across the different sites is given in Figure ii. For the height growth, we had 3057 observations, while there were 5514 survival assessments. Note that no height growth data is given for SAPET and SICH, as we omitted these sites in the growth rate analysis.

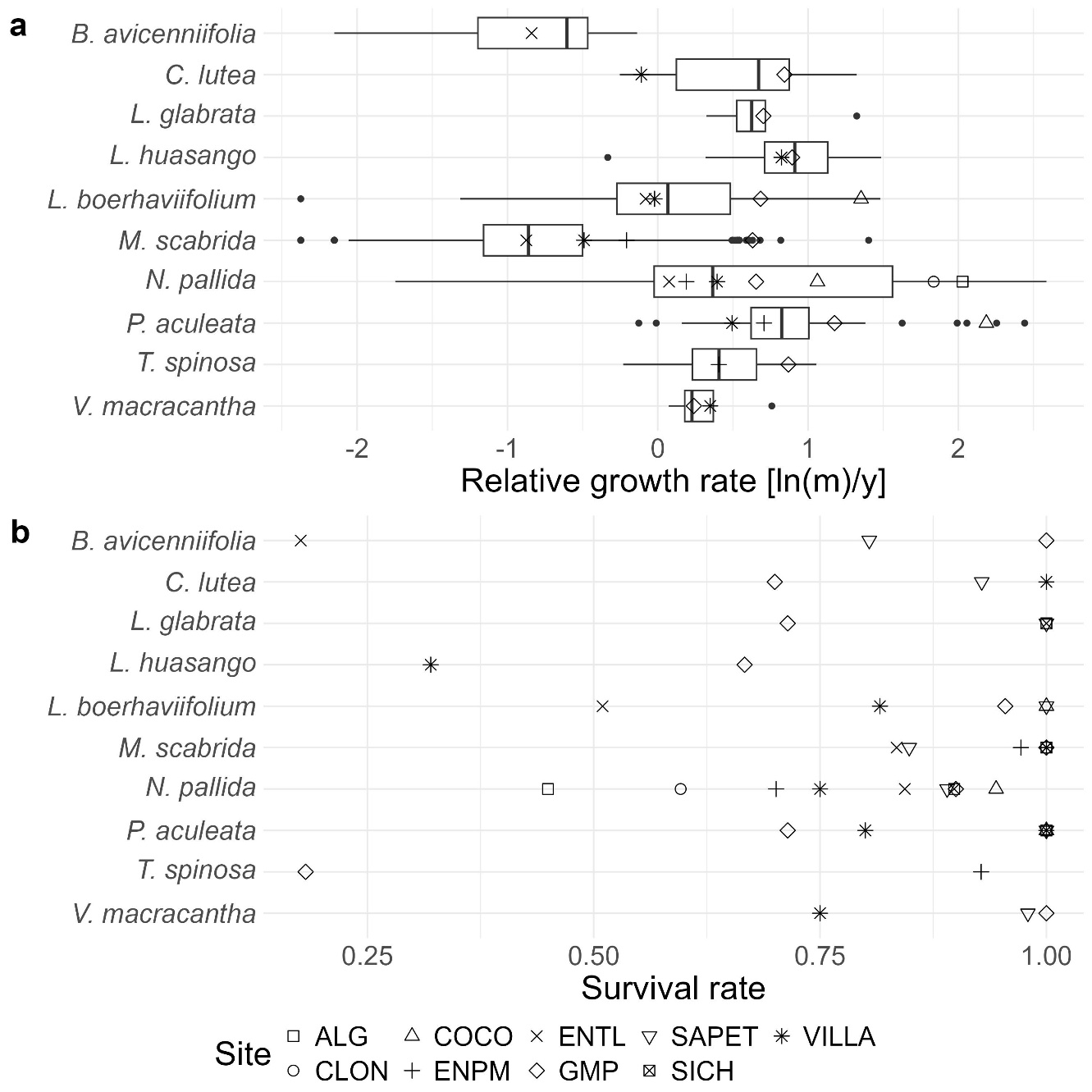

Figure ii: Overview of relative growth (a) and survival (b) rates across the nine sites for the ten species studied (i.e. Beautempsia avicenniifolia, Cordia lutea, Libidibia glabrata Loxopterygium huasango, Lycium boerhaviifolium, Morisonia scabrida, Neltuma pallida, Parkinsonia aculeata, Tara spinosa and Vachellia macracantha).

### Supplement S4: AGE model survival

As we could not include both age and the random effect of the site in our survival models, we included here the fixed effect of age for the model focusing on the interactive effect between resource-use strategy and potential evapotranspiration, omitting the random effect of site (Table vi). Note that the fixed effect of age becomes highly significant (AGE -0.04 p < 0.001) and increases the marginal R² to 11.7%, while the interaction between resource-use strategy and potential evapotranspiration is no longer statistically significant, reinforcing our choice not to place too much importance on the survival models.

*Table vi: Parameter estimates (β) and 95%-confidence intervals (CI) for the generalised linear mixed models (GLMMs) examining survival rates focusing on the interaction between resource-use strategy and water availability and incorporating the effect of age, thus excluding the random effect of site. Predictor variables include nitrogen fixation (Nfix), resource-use strategy (PC_resource-use_), size variation (PC_size_), potential evapotranspiration (PET [mm/year]), organic matter content (OM [%]), mean temperature in the wettest quarter (BIO8 [°C]) and site age. Low values of PC_resource-use_* *indicate acquisitive species, while higher values correspond to more conservative species. Significance of the parameters is indicated with * p<0.05, ** p<0.01 and *** p<0.001. Goodness-of-fit is given as both marginal (R²m) and conditional (R²c) coefficients of determination.*

| **Driver** |  | **Survival rate** | |
| --- | --- | --- | --- |
|  |  | β | CI |
| Nfix |  | 0.52 | [0.11, 0.93] |
| PC_resource-use_ |  | -0.12 | [-0.26, 0.02] |
| PC_size_ |  | -0.15 | [-0.34, 0.04] |
| PET |  | -0.02 | [-0.07, 0.02] |
| OM |  | -0.02 | [-0.05, 0.01] |
| BIO8 |  | 0.00 | [-0.07, 0.06] |
| AGE |  | -0.04*** | [-0.05, -0.04] |
| PC_resource-use_*PET | | -0.04 | [-0.06, -0.01] |
| PC_resource-use_*OM |  | 0.05 | [0.02, 0.07] |
| R²_m_ (%) |  | 11.7 |  |
| R²_c_ (%) |  | 23.8 |  |

### Supplement S5: BIO8 interaction model

Due to high VIFs, we were not able to incorporate the interaction between resource-use strategy and BIO8 in our main model. We therefore tested a second model where the interaction between PC_resource-use_ and PET was omitted and replaced by that between PC_resource-use_ and BIO8 (Table vii). In this supplement, we give the parameter estimates and 95%-confidence intervals for all variables, while the main text only focuses on the significant interactions.

Table vii: Parameter estimates (β) and 95%-confidence intervals (CI) for the generalised linear mixed models (GLMMs) examining height growth and survival rates focusing on the interaction between resource-use strategy and temperature in the wettest quarter. Predictor variables include nitrogen fixation (Nfix), resource-use strategy (PC_resource-use_), size variation (PC_size_), potential evapotranspiration (PET [mm/year]), organic matter content (OM [%]), mean temperature in the wettest quarter (BIO8 [°C]) and site age. Low values of PC_resource-use_ indicate acquisitive species, while higher values correspond to more conservative species. Significance of the parameters is indicated with * p<0.05, ** p<0.01 and *** p<0.001. Goodness-of-fit is given as both marginal (R²_m_) and conditional (R²c) coefficients of determination.

| **Driver** |  | **Height growth rate** | |  | **Survival rate** | |
| --- | --- | --- | --- | --- | --- | --- |
|  |  | β | CI |  | β | CI |
| Nfix |  | -0.16 | [-0.35, 0.02] |  | 0.47 | [0.06, 0.89] |
| PCresource-use |  | -0.42*** | [-0.49, -0.36] |  | -0.15 | [-0.30, 0.00] |
| PCsize |  | 0.16 | [0.08, 0.24] |  | -0.12 | [-0.31, 0.08] |
| PET |  | -0.32* | [-0.41, -0.24] |  | 0.05 | [-0.11, 0.21] |
| OM |  | 0.10* | [0.06, 0.15] |  | -0.05 | [-0.08, -0.01] |
| BIO8 |  | 0.49* | [0.39, 0.59] |  | -0.25 | [-0.41, -0.08] |
| AGE |  | 0.04* | [0.04, 0.05] |  | NA | NA |
| PCresource-use*BIO8 | | -0.16*** | [-0.18, -0.14] |  | -0.06 | [-0.10, -0.01] |
| PCresource-use*OM |  | 0.09*** | [0.07, 0.12] |  | 0.07* | [0.04, 0.09] |
| R²m (%) |  | 75.3 |  |  | 4.7 |  |
| R²c (%) |  | 82.3 |  |  | 24.7 |  |

### Supplement S6: Original models full tables

Here, we provide a more complete version of Table 1, which shows only the parameter estimate (β) and confidence interval (CI) for the model focusing on the interaction between resource-use strategy and potential evapotranspiration (Table viii). We added the degrees of freedom (DF) and t-values to the height growth rate model, while the z-values are given for the survival rate model.

Table viii: Parameter estimates (β) and 95%-confidence intervals (CI) for the generalised linear mixed models (GLMMs) examining height growth and survival rates focusing on the interaction between resource-use strategy and water availability. Predictor variables include nitrogen fixation (Nfix), resource-use strategy (PC_resource-use_), size variation (PC_size_), potential evapotranspiration (PET [mm/year]), organic matter content (OM [%]), mean temperature in the wettest quarter (BIO8 [°C]) and site age. Low values of PC_resource-use_ indicate acquisitive species, while higher values correspond to more conservative species. Significance of the parameters is indicated with * p<0.05, ** p<0.01 and *** p<0.001. Goodness-of-fit is given as both marginal (R²_m_) and conditional (R²c) coefficients of determination.

| **Driver** |  | **Height growth rate** | | | |  | **Survival rate** | | |
| --- | --- | --- | --- | --- | --- | --- | --- | --- | --- |
|  |  | β | CI | DF | t-value |  | β | CI | z-value |
| Nfix |  | -0.17 | [-0.36, 0.02] | 7.98 | -0.91 |  | 0.48 | [0.06, 0.90] | 1.15 |
| PC_economics_ |  | -0.29** | [-0.35, -0.23] | 5.94 | -4.60 |  | -0.11 | [-0.25, 0.04] | -0.76 |
| PC_size_ |  | 0.16 | [0.07, 0.24] | 7.27 | 1.91 |  | -0.11 | [-0.30, 0.09] | -0.58 |
| PET |  | -0.31* | [-0.40, -0.23] | 3.41 | -3.71 |  | 0.05 | [-0.11, 0.21] | 0.29 |
| OM |  | 0.10* | [0.06, 0.14] | 28.15 | 2.36 |  | -0.04 | [-0.08, -0.01] | -1.26 |
| BIO8 |  | 0.48* | [0.38, 0.57] | 3.43 | 4.81 |  | -0.25 | [-0.41, -0.09] | -1.57 |
| AGE |  | 0.04* | [0.04, 0.05] | 3.09 | 5.21 |  | NA | NA | NA |
| PC_economics_*PET | | -0.09*** | [-0.11, -0.08] | 1501.83 | -7.65 |  | -0.03 | [-0.06, 0.00] | -1.046 |
| PC_economics_*OM |  | 0.10*** | [0.07, 0.12] | 295.71 | 4.14 |  | 0.06* | [0.03, 0.09] | 2.24 |
| R²_m_ (%) |  | 75.3 |  |  |  |  | 4.6 |  |  |
| R²_c_ (%) |  | 82.3 |  |  |  |  | 24.6 |  |  |
